# Mitochondrial phosphoproteomes are functionally specialized across tissues

**DOI:** 10.1101/2022.03.23.485457

**Authors:** Fynn M. Hansen, Laura S. Kremer, Ozge Karayel, Inge Kühl, Isabell Bludau, Nils-Göran Larsson, Matthias Mann

## Abstract

Mitochondria are essential organelles involved in critical biological processes such as energy metabolism and cell survival. Their dysfunction is linked to numerous human pathologies that often manifest in a tissue-specific manner. Accordingly, mitochondrial fitness depends on versatile proteomes specialized to meet diverse tissue-specific requirements. Furthermore, increasing evidence suggests that phosphorylation may also play an important role in regulating tissue-specific mitochondrial functions and pathophysiology. We hypothesized that recent advances in mass spectrometry (MS)-based proteomics would now enable in-depth measurement to quantitatively profile mitochondrial proteomes along with their matching phosphoproteomes across tissues. We isolated mitochondria from mouse heart, skeletal muscle, brown adipose tissue, kidney, liver, brain, and spleen by differential centrifugation followed by separation on Percoll gradients and high-resolution MS analysis of the proteomes and phosphoproteomes. This in-depth map substantially quantifies known and predicted mitochondrial proteins and provides a resource of core and tissue modulated mitochondrial proteins (mitophos.de). We also uncover tissue-specific repertoires of dozens of kinases and phosphatases. Predicting kinase substrate associations for different mitochondrial compartments indicates tissue-specific regulation at the phosphoproteome level. Illustrating the functional value of our resource, we reproduce mitochondrial phosphorylation events on DRP1 responsible for its mitochondrial recruitment and fission initiation and describe phosphorylation clusters on MIGA2 linked to mitochondrial fusion.

## Introduction

Mitochondria are double-membrane-bound organelles with an essential role in homeostasis of eukaryotic cells. They are often referred to as the “powerhouse of the cell” due to their prominent function in bioenergetics. Among many other processes, they are also involved in several biosynthetic processes such as balancing redox systems, the regulation of metabolic by-products like reactive oxygen species (ROS) (Spinelli and Haigis, 2018) and hold a central role in cell death (Bock and Tait, 2020). The function and stability of mitochondria depend on their intrinsic bioenergetics regulation and finely orchestrated interaction with the cellular microenvironment. Energy conversion via the oxidative phosphorylation system (OXPHOS) plays an essential role in harvesting energy from ingested nutrients. Moreover, the morphology of mitochondria within an eukaryotic cell is actively regulated by fusion and fission events which dynamically modulate their number, size, and localization (Liesa et al., 2009). Regulation of mitochondrial dynamics also affects the interplay of mitochondria with other cellular structures, such as the cytoskeleton for active regulation of their localization (Moore and Holzbaur, 2018), and organelles like the endoplasmic reticulum (ER) and lipid droplets to regulate many physiological processes such as energy metabolism and ion buffering. The mitochondria-associated membrane (MAM), which is the contact site of the outer mitochondrial membrane with the ER, comprises a unique set of proteins mediating this interaction and fine-tune mitochondrial functions with the cellular microenvironment (Kwak et al., 2020; Nunnari and Suomalainen, 2012). Dysregulation of any of these intricate processes can lead to severe mitochondrial dysfunctions and diseases, including neurodegenerative diseases, cardiovascular disorders, myopathies, obesity, and cancers, which can manifest in a cell type- and tissue-specific manner (Suomalainen and Battersby, 2018).

The fitness of mitochondria depends on the production and maintenance of functional as well as versatile proteomes specialized to carry out a variety of functions within the eukaryotic metabolism and meet diverse cellular and tissue-specific requirements (Kuznetsov et al., 2009). The mitochondrial proteome includes over a thousand proteins (see below), but only a small fraction of 13 proteins are encoded on the circular mitochondrial DNA molecule (Anderson et al., 1981). Thus, the majority of mitochondrial proteins are encoded by the nuclear genome, synthesized outside of mitochondria and subsequently imported into the organelle, implying that mechanisms controlling mitochondrial protein quality (e.g., correct protein folding and import) are essential for health and integrity of mitochondria (Jadiya and Tomar, 2020). Furthermore, investigations of mitochondrial dynamics and functional plasticity have revealed regulatory roles for post-translational modifications (PTMs), including phosphorylation (Niemi and Pagliarini, 2021). Studies have shown that phosphorylation of several mitochondrial proteins is involved in the regulation of central processes such as metabolic function, for instance through phosphorylation of the E1alpha subunit of PDH (Patel et al., 2014), mitophagy (Kolitsida et al., 2019) and fission (Cribbs and Strack, 2007; Ducommun et al., 2015; Lewis et al., 2018; Taguchi et al., 2007; Toyama et al., 2016). Thus, deregulation of protein phosphorylation might be an important underlying feature of mitochondrial physiology and pathophysiology. Currently, there is a significant knowledge gap of the mitochondrial variable proteomic composition and to what extend it is phosphorylated in a tissue-specific manner and how post-translational regulation influences organelle function. A detailed understanding of the functional specialization of mitochondria at the protein and phosphorylation levels is needed to elucidate the contribution of mitochondria to health and disease.

Large-scale mass spectrometry (MS)-based quantitative proteomics studies from our and other groups have already shed light on the proteomic composition of mitochondria of various mammalian tissues and cell types, mostly highlighting that the majority of proteins are shared between mitochondria of different tissues (Forner et al., 2006; Mootha et al., 2003; Pagliarini et al., 2008). The breadth and depth of such studies has been largely driven by technological advances in the field in combination with improvements of mitochondria isolation procedures, such as differential centrifugation (DC), DC in conjunction with ultracentrifugation on e.g. Percoll gradients, magnetic bead-assisted methods (MACS) (Kappler et al., 2016) or MitoTags (Bayraktar et al., 2019). Efforts in defining the mitochondrial proteome lead to databases like MitoCarta2.0 and IMPI (http://impi.mrc-mbu.cam.ac.uk/), both integrated in Mitominer4.0 (Smith and Robinson, 2019). A recent quantitative and high confidence proteome of human mitochondria identified 1134 different proteins that vary over six orders in magnitude in abundance (Morgenstern et al., 2021). Similarly, efforts have been undertaken to map the mitochondrial phosphoproteome, and identified dozens to hundreds of phosphorylation sites on mitochondrial proteins (listed in Supplementary Table 1). However, there has been a dramatic improvement in the technology of phosphoproteomics workflows during the last years, leading to the routine identification and quantification of thousands of phosphorylation sites in cell culture and in vivo systems (Bekker-Jensen et al., 2020; Humphrey et al., 2018) which had not been available in earlier studies. Furthermore, comparisons between mitochondrial phosphoproteomes have been difficult because only a single or a few tissues were analyzed. This complicates the combination and comparison of data sets across studies to obtain a clear view of mitochondrial diversity on proteome and phosphoproteome levels. Thus, a concerted effort is needed to systematically and quantitatively profile mitochondrial proteomes together with their matching phosphoproteomes from the same biological source. This would further help to investigate the dynamic composition of mitochondria and help identify the tissue-specific repertoire of mitochondria-resident kinases and phosphatases and their substrate associations.

Here we performed a systematic analysis of the mitochondrial composition at the level of proteins, major functional entities, and phosphorylation in seven mouse tissues – brain, brown adipose tissue (BAT), heart, kidney, liver, skeletal muscle (SKM), and spleen. Our study employs state of the art MS-based proteomics technology to systematically map divergent composition and phosphorylation of mitochondria between tissues and provides functionally valuable insights into their proteome and post-translational regulations. This study contributes to our understanding of tissue-specific mitochondrial processes controlled by protein abundance and phosphorylation and helps to manipulate these in health and disease. Our mitochondrial (phopho)proteomes are composed into an extensive resource and made freely accessible via mitophos.de.

## Results

### Comprehensive mitochondria proteome coverage across various mouse tissues

To advance our understanding of tissue-specific functional specializations of mitochondria at the protein level, we set out to characterized proteomes of mitochondria collected from various mouse tissues by LC-MS/MS analysis. To this end, we first isolated mitochondria from seven tissues – brain, brown adipose tissue (BAT), heart, kidney, liver, skeletal muscle (SKM), spleen - from six 18-21 weeks old C57BL/6N mice (three females and three males). Mouse tissues were homogenized with a Dounce homogenizer, and crude mitochondria were isolated via differential centrifugation and subsequently purified on a Percoll density gradient to obtain ultra-pure mitochondria isolates (Kuhl et al., 2017). Importantly, this procedure was shown to efficiently exclude contaminations from other cellular compartments (Wieckowski et al., 2009). Proteomes of these ultra-pure mitochondrial samples were acquired by a state-of-the-art proteomics workflow (Figure 1A), allowing the robust identification and quantification (coefficient of variation (CV) < 20%) of proteins that covered a dynamic range of more than five orders of magnitude (Supplementary File 1).

**Figure 1.**
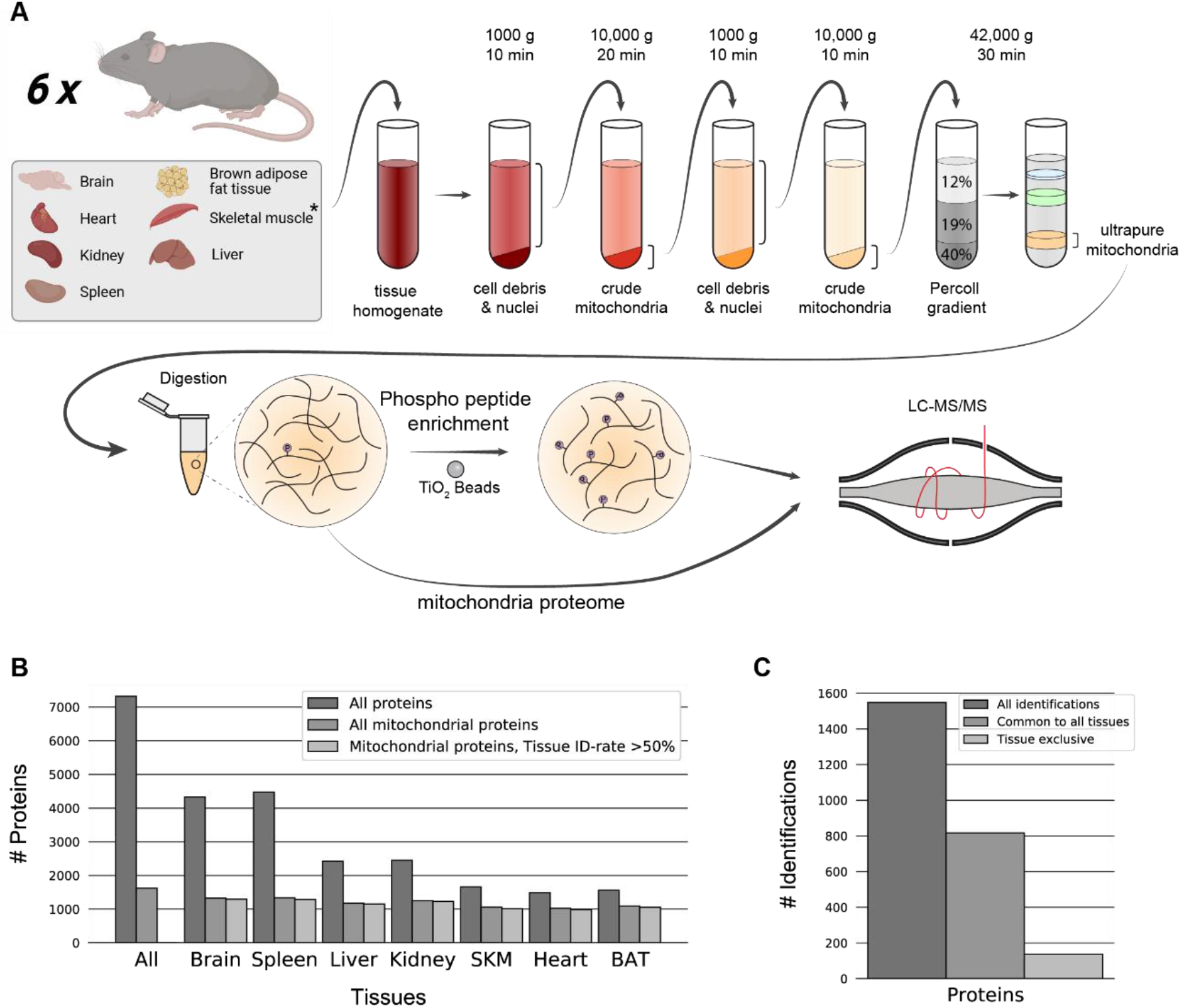
Mitochondrial proteome and phosphoproteome preparation. (A) Workflow of tissue preparations for mitochondrial proteome and phosphoproteome enrichment, and LC-MS/MS analysis (n=6). Tissues were first homogenized (* for skeletal muscle, see Methods), crude mitochondria were isolated and ultrapure mitochondria were obtained using a Percoll gradient. Proteins were digested and prepared for phosphoproteome analysis via TiO2 enrichment or subjected to LC-MS/MS. (B) Distinct protein identification across biological replicates (n=6). Annotation of proteins as mitochondrial is based on MitoCarta3.0 and the IMPI database. (C) Mitochondrial protein numbers after filtering for mitochondrial proteins identified in at least 50% of biological replicates (n=6) in at least one tissue. Skeletal muscle (SKM), brown adipose tissue (BAT). See also Figure 1 – Source Data 1.

In total, we identified over 7000 proteins of which 1620 were annotated as mitochondrial proteins using MitoCarta3.0 (Rath et al., 2021) and the IMPI (http://impi.mrc-mbu.cam.ac.uk/) database. This essentially covers (92%) the mitochondrial proteome by the measure of MitoCarta3.0 and even in the IMPI database, which also includes predicted mitochondrial proteins, we still identified 62% (Figure 1B). For further analysis, we filtered for proteins identified in more than half of the biological replicates in at least one tissue, which resulted in 1548 mitochondrial proteins. This still represents over 90% of MitoCarta3.0 and 59% of the IMPI databases and highlights the deep and reproducible mitochondrial proteome coverage of this study (Figure 1C). Interestingly, half of these mitochondrial proteins were identified across all tissues while only 9% were exclusively detected in one specific tissue (Figure 1C), confirming previous reports on mitochondrial proteomes by us and others (Calvo and Mootha, 2010; Forner *et al*., 2006; Johnson et al., 2007; Mootha *et al*., 2003). Of these, almost half (65 proteins) were both reproducibly identified and not in the lowest 20% of ranked abundances (Supplementary File 2), making them clear candidates for tissue specific mitochondrial proteins. A similar proportion of the mitochondrial proteome was also exclusive to two or more tissues by the same criterion.

Mitochondrial enrichment efficiency can be determined by the proportion of summed signal intensity for mitochondrial proteins in relation to all identified proteins in measured samples (Williams et al., 2018). Applying this strategy, we determined the proportions to be very high (>95%) for liver, kidney, SKM, heart and BAT, but lower for brain (65%) and spleen (45%) (Figure 2A). These trends are consistent with the established literature (Fecher et al., 2019; McLaughlin et al., 2020; Roberts et al., 2021; Williams *et al*., 2018) and can likely be explained by the high tissue heterogeneity of brain (Fecher *et al*., 2019; Menacho and Prigione, 2020) and spleen. Indeed, spleen tissue consists of various types of immune cells (Lewis et al., 2019), which might impede high purity enrichment of its organelles. For brain, contaminations by synaptosomes, which themselves contain neuronal mitochondria, have frequently been observed (Mootha *et al*., 2003). Yet, correlation of both mitochondrial and all identified proteins between biological replicates yielded Pearson correlation coefficients higher than 95% in all tissues (Figure 2B and Figure 2- figure supplement 1A). Principle component analysis (PCA) further shows that biological replicates cluster together and underlines functional similarities between tissues such as heart and SKM as well as tissue related diversity of mitochondria proteomes (Figure 2C and Figure 2-figure supplement 1B).

**Figure 2.**
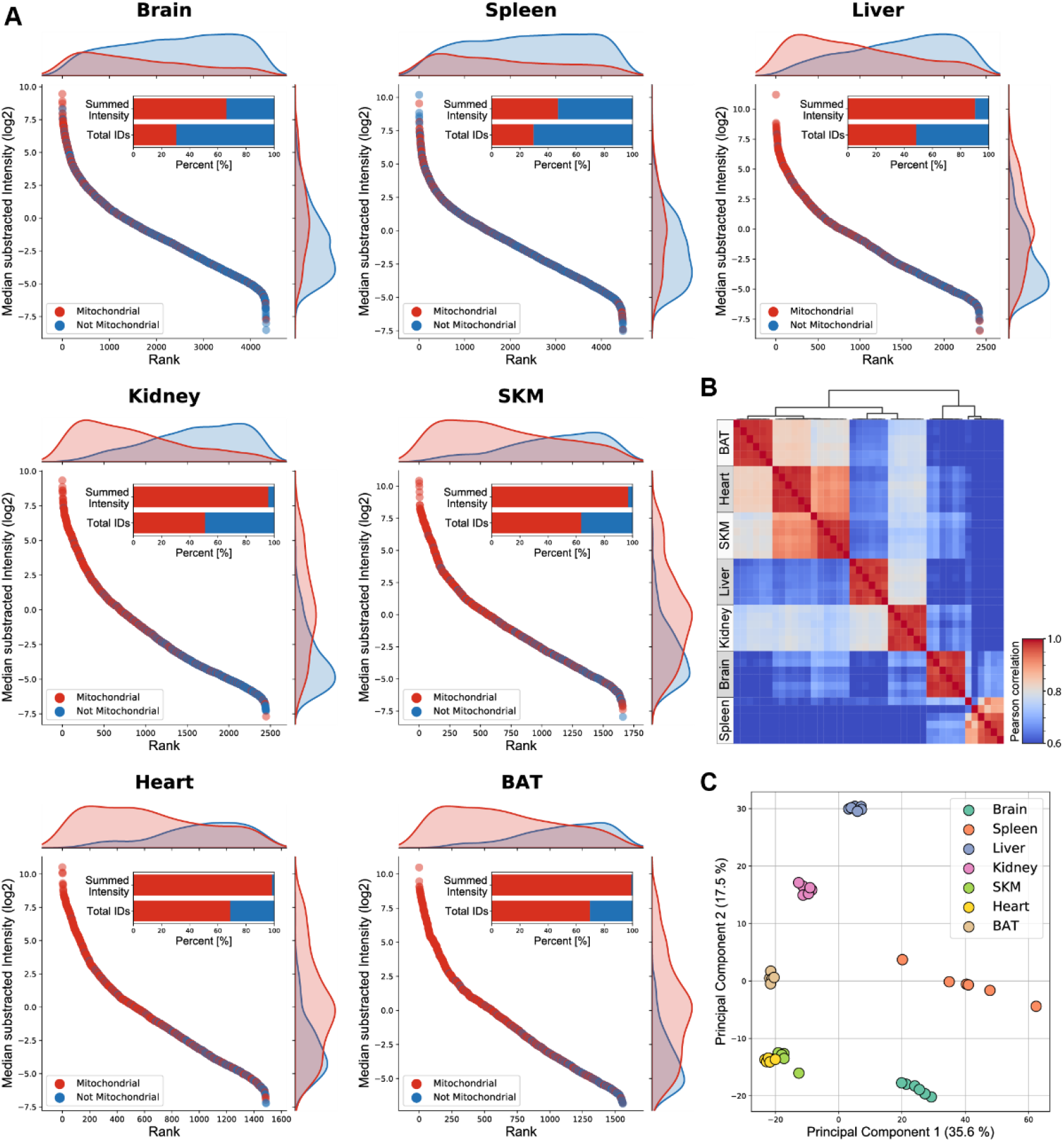
Mitochondria enriched samples show tissue-specific clustering. (A) Mitochondrial (red) and not mitochondrial (blue) proteins identified, based on the MitoCarta3.0 and IMPI database, are ranked by their intensity for each individual tissue. Histograms on the top and right display the distribution of mitochondrial (red) and not mitochondrial (blue) proteins along the rank and the Intensity axis, respectively. The percentage of all identified mitochondrial (red) and not mitochondrial (blue) proteins and their summed intensities are displayed in bar graphs. (B) Heatmap showing Pearson correlation coefficient for biological replicates (n=6) for mitochondrial proteins of all mitochondrial enriched samples. (C) Principal component analysis of mitochondrial proteins of all acquired biological replicates (n=6). Skeletal muscle (SKM), brown adipose tissue (BAT). See also Figure 2 – Source Data 1.

### Mitochondrial proteome composition reveals tissue-specific functions

To gain further insights into tissue-specific functional differences in mitochondria, we investigated differences in the composition of mitochondrial proteomes across tissues. First, we focused on the oxidative phosphorylation (OXPHOS) system, which is essential for production of the energy-rich metabolite ATP and other processes like free radical generation and apoptosis (Huttemann et al., 2007). We found that proteins of the electron transport chain (Complex I – Complex IV) and ATP-synthase (Complex V) displayed high abundances in heart and SKM tissues, supporting the physiological requirement of high ATP levels in both muscular tissues to sustain processes like muscle contraction (Ferreira et al., 2010; Ventura-Clapier et al., 2011) (Figure 3A). Conversely, levels of Complex V proteins were substantially lower in mitochondria from BAT compared to all other measured tissues (Figure 3A), which agrees with the specialized function of BAT in non-shivering thermogenesis (Jastroch et al., 2010; Kajimura and Saito, 2014; Oelkrug et al., 2015). This was further supported by the high abundance of the uncoupling protein 1 (UCP1) in our proteome measurements, the key mediator of the heat-generating proton leak in the mitochondria of BAT (Figure 3B).

**Figure 3.**
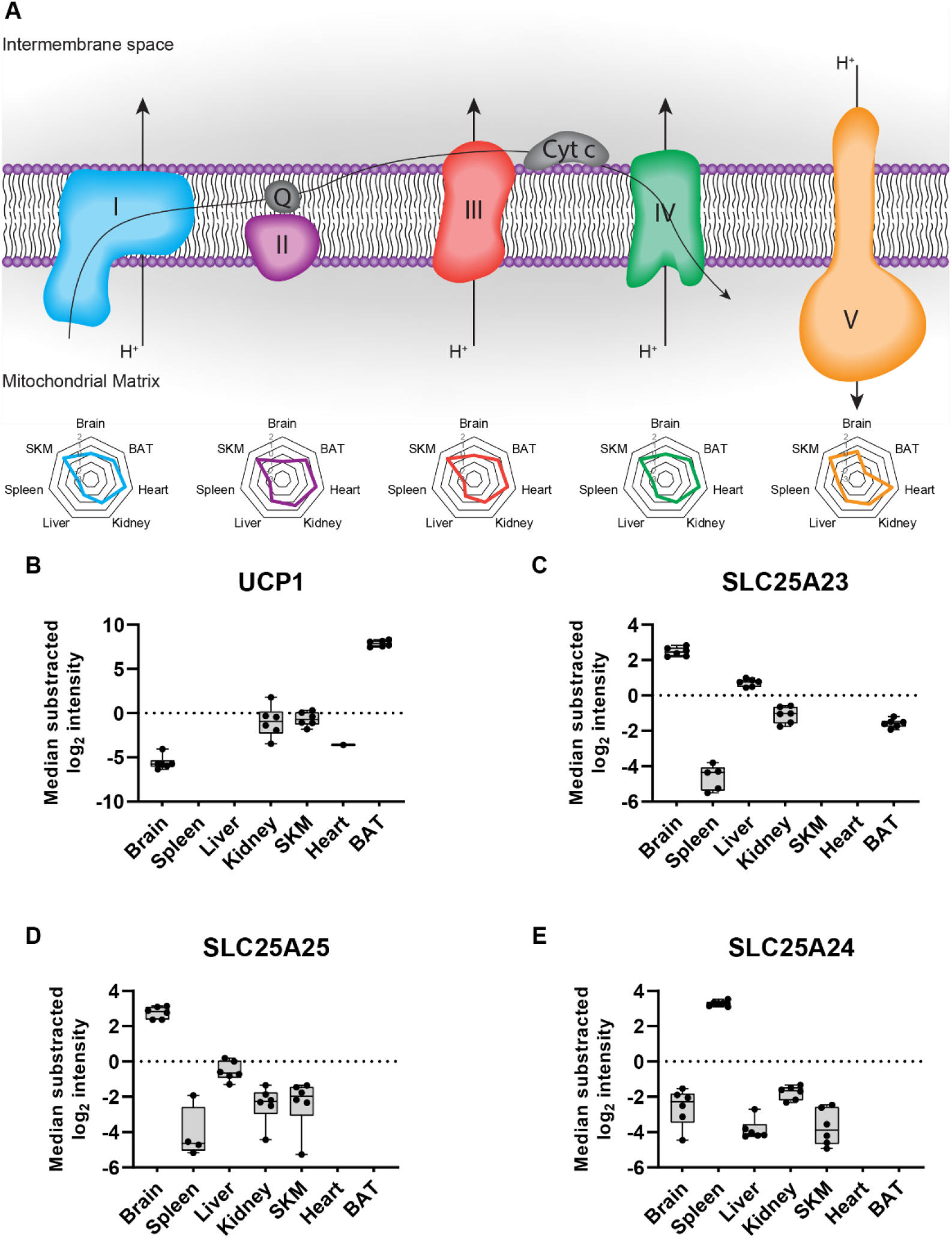
Tissue-specific protein contribution of SLC25 proteins the mitochondrial proteome. (A) Representation of the oxidative phosphorylation (OXPHOS) system including from left to right the electron transport chain (Complex I (blue), Complex II (violet), Complex III (red), Complex IV (green)) and ATP synthase (Complex V (yellow)). Radar plots show the relative contribution of the corresponding complex across the analyzed tissues to the overall mitochondrial composition (see methods for detailed description). (B) Normalized intensity (median of all log2 transformed mitochondrial proteins of a sample was subtracted from all log2 protein intensities of that sample) of the uncoupling protein 1 (UCP1), (C) SLC25A23, (D) SLC25A25, and (E) SLC25A24 across all analyzed tissues (black dots indicate individual identifications). Protein abundance differences in relation to the reference tissue (BAT in B, Brain in C and D and spleen in E) were significant (p-value<0.0001) by one-way ANOVA analysis (Figure 3 - Source Data 1). Data in this figure is based on the analysis of six replicates (n=6) for each tissue. Skeletal muscle (SKM), brown adipose tissue (BAT). See also Figure 3 – Source Data 1.

In contrast to the well described UCP1, several other members of the SLC25 family remain uncharacterized (Ruprecht and Kunji, 2020). In our dataset, we identified a total of 47 members of the SLC25 family (Kunji et al., 2020). We found that levels of SLC25A23 and SLC25A25, two ATP-Mg^2+^/P_i_ carrier paralogues, (del Arco and Satrustegui, 2004; Fiermonte et al., 2004), displayed high abundance in mitochondria of brain tissue (Figure 3C and Figure 3D). Interestingly, knockout of SLC25A23 was shown to increase neuronal vulnerability (Rueda et al., 2015), which corroborates our observation and suggests an important role of SLC25A25 in this tissue type. We also quantified a third paralogue, SLC25A24, which showed a higher abundance in spleen tissue. Notably, the spleen harbors a large pool of B-cells and reduced SLC25A24 levels were previously linked to B-cell malignancies (Sandhu et al., 2013) (Figure 3E). Although these three ATP-Mg^2+^/P_i_ carriers were not detected in mitochondria from heart tissue, we identified another class of ATP carriers, including SLC25A4 or SLC25A31, which both showed increased abundance in heart compared to all other measured tissues. Such differences between mitochondrial proteome compositions are easily retrieved from our dataset, facilitating a better understanding of mitochondrial plasticity across tissues.

### Tissue-specific function of mitochondria-associated proteins

The identification of key proteins mediating the crosstalk of mitochondria with their cellular environment is crucial to better understand their tissue-specific regulation and this concept has already attracted considerable interest in recent years (Montes de Oca Balderas, 2021). Although the characterization of local proteomes of organellar contact sites usually require special centrifugation-based isolation methods, we anticipated that a considerable fraction of mitochondria-associated proteins would also enrich along with mitochondria in our samples. We first performed an annotation term enrichment analysis of all identified proteins. While we observed significant enrichment of several mitochondria related terms in all tissues as expected, the non-mitochondrial protein pool was largely enriched for terms related to the tissue of origin (Figure 4 – supplement 1, Figure 4 - Source Data 1). For instance, terms like ‘epoxygenase P450 pathway’ in liver, ‘positive regulation of B cell activation’ in spleen or ‘positive regulation of excitatory postsynaptic potential’ in brain tissue highlight tissue-specific functions.

**Figure 4.**
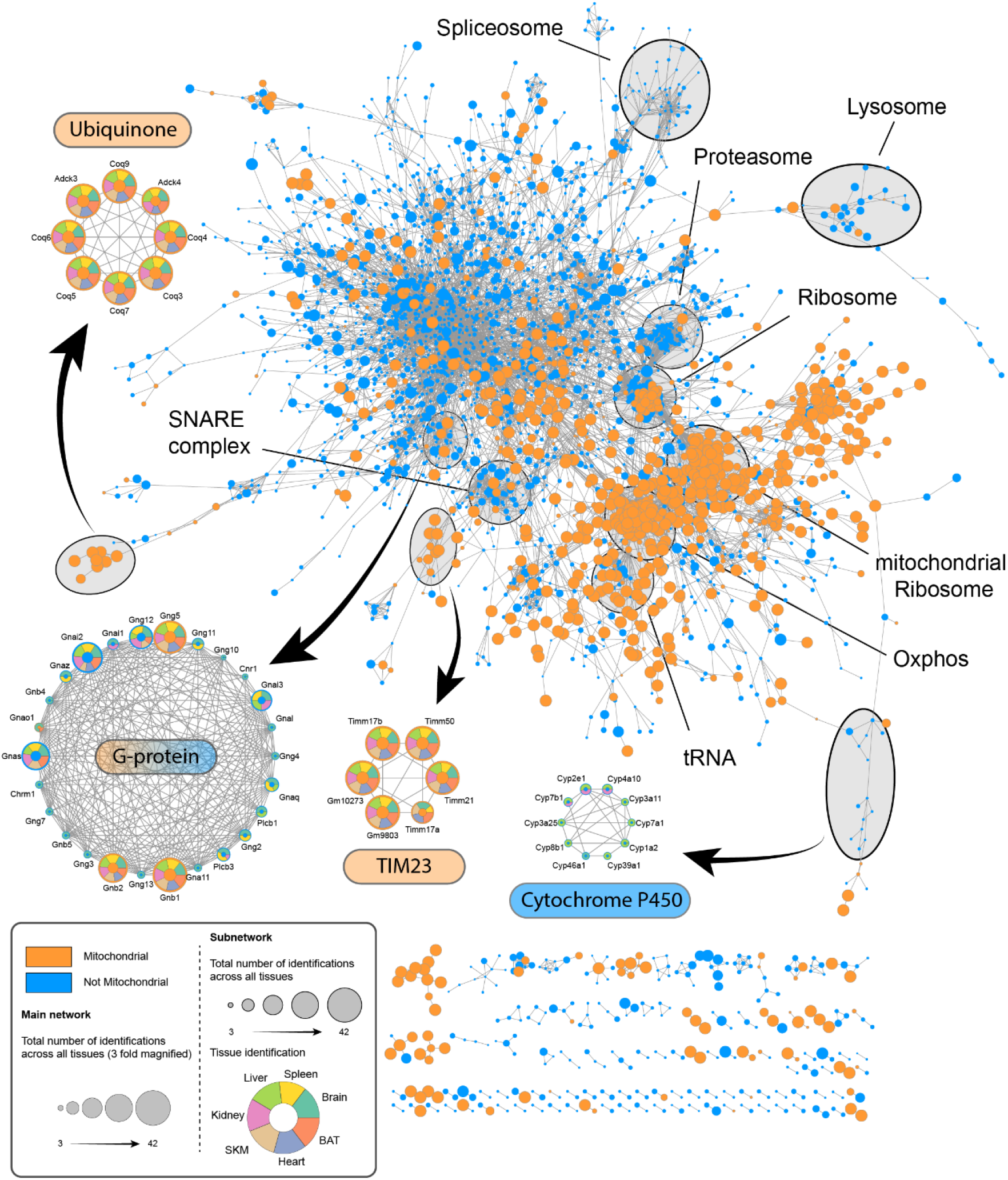
Proteome of mitochondria enriched samples displays tissue-specific complexes. Cytoscape network analysis of reproducibly (> 50% identification rate in at least one tissue, six replicates were analyzed (n=6)) identified proteins of all mitochondria enriched samples. The main network depicts mitochondrial (orange) and non- mitochondrial (blue) proteins with at least one edge (String score >0.95). The size of individual nodes represents the number of identifications ranging from 3 (small circle) to 42 (big circle). Subnetworks display mitochondrial (orange) and non-mitochondrial (blue) proteins and tissues in which they were identified (dark green – brain; yellow – spleen; light green – liver; pink – kidney; ochre – SKM; blue – heart; orange – BAT). Skeletal muscle (SKM), brown adipose tissue (BAT). See also Figure 4 – Source Data 1.

Next, we performed a network analysis to evaluate the nature and quality of co-enriched proteins, more specifically whether non-mitochondrial proteins identified in these samples are associating proteins with functional roles or biological contaminations that are likely tissue-specific and high abundant. This analysis revealed several clusters of known mitochondrial complexes such as the Tim23 complex or processes like the ubiquinone biosynthetic process, whose members were robustly identified in all tissues, and several tissue-specific clusters including both mitochondrial and non-mitochondrial proteins (Figure 4). For instance, in line with terms enriched for co-sedimenting proteins in liver tissue, we identified a cluster of cytochrome P450 superfamily members exclusively in liver tissue (Figure 4). Intriguingly, CYP2E1 and CYP1A2, two specific members of this family, were previously shown to be targeted to the mitochondria (Avadhani et al., 2011; Genter et al., 2006; Robin et al., 2001), and we now find evidence of the enrichment of many more family members in this tissue. Another identified cluster consisted of G proteins, some of which were identified throughout all tissues and previously annotated as mitochondrial (e.g. GNB1, GNB2 and GNG5) or shown to localize to mitochondria (GNAI2) (Beninca et al., 2014). Interestingly, like the majority of the proteins in the G protein cluster, many G protein coupled receptors (GPCR) were exclusively identified in brain tissue. A prominent member of these brain specific GPCRs is CNR1, which was reported to localize to mitochondria where it plays an important role in the regulation of memory processes through the modulation of the mitochondrial energy metabolism (Hebert-Chatelain et al., 2016). These examples highlight the value of our dataset for uncovering novel mitochondrial and mitochondria-associated proteins in an unbiased way.

### Mitochondrial kinases and phosphatases show tissue specificity

Post-translational modification of proteins, specifically phosphorylation, plays a crucial role in the orchestration of mitochondrial protein function (Niemi and Pagliarini, 2021). However, almost no mitochondrial kinases with mitochondrial targeting sequences have been consistently reported and most kinases shown to associate with mitochondria have been found on or interact with the outer membrane (Kotrasova et al., 2021). Given our deep mitochondrial proteomes, we investigated relative abundances of kinases and phosphatases, which are annotated as or suggested to be mitochondrial, across mouse tissues.

Firstly, we observed clear differences in abundances of identified mitochondrial kinases and phosphatases, including well described matrix kinases and phosphatases, between tissues (Figure 5A). For instance, PDK1, PDK2, and PDK4 contributed preferentially to the composition of heart, SKM and BAT mitochondria, while the PDK3 was more abundant in brain, spleen, and kidney mitochondria. These tissue-related differences are in line with earlier reports and suggest a specialized function of PDK3, which may originate in its insensitivity to pyruvate inhibition (Klyuyeva et al., 2019; Sadana et al., 2018). Similarly, levels of the heterodimeric pyruvate dehydrogenase phosphatase consisting of PDP1 and PDPr were elevated in brain, SKM, and heart compared to the remaining tissues, whereas PDP2 contributes more to the composition of liver, kidney, and BAT mitochondria. While our study confirmed previous reports on the differential expressions of these proteins in a tissue-specific manner (Huang et al., 1998; Huang et al., 2003), it also provided quantitative data to assess the magnitude of these differences. Together, our results imply a tailored regulation of kinase and phosphatase abundances across tissues to modulate the mitochondrial phosphoproteome.

**Figure 5.**
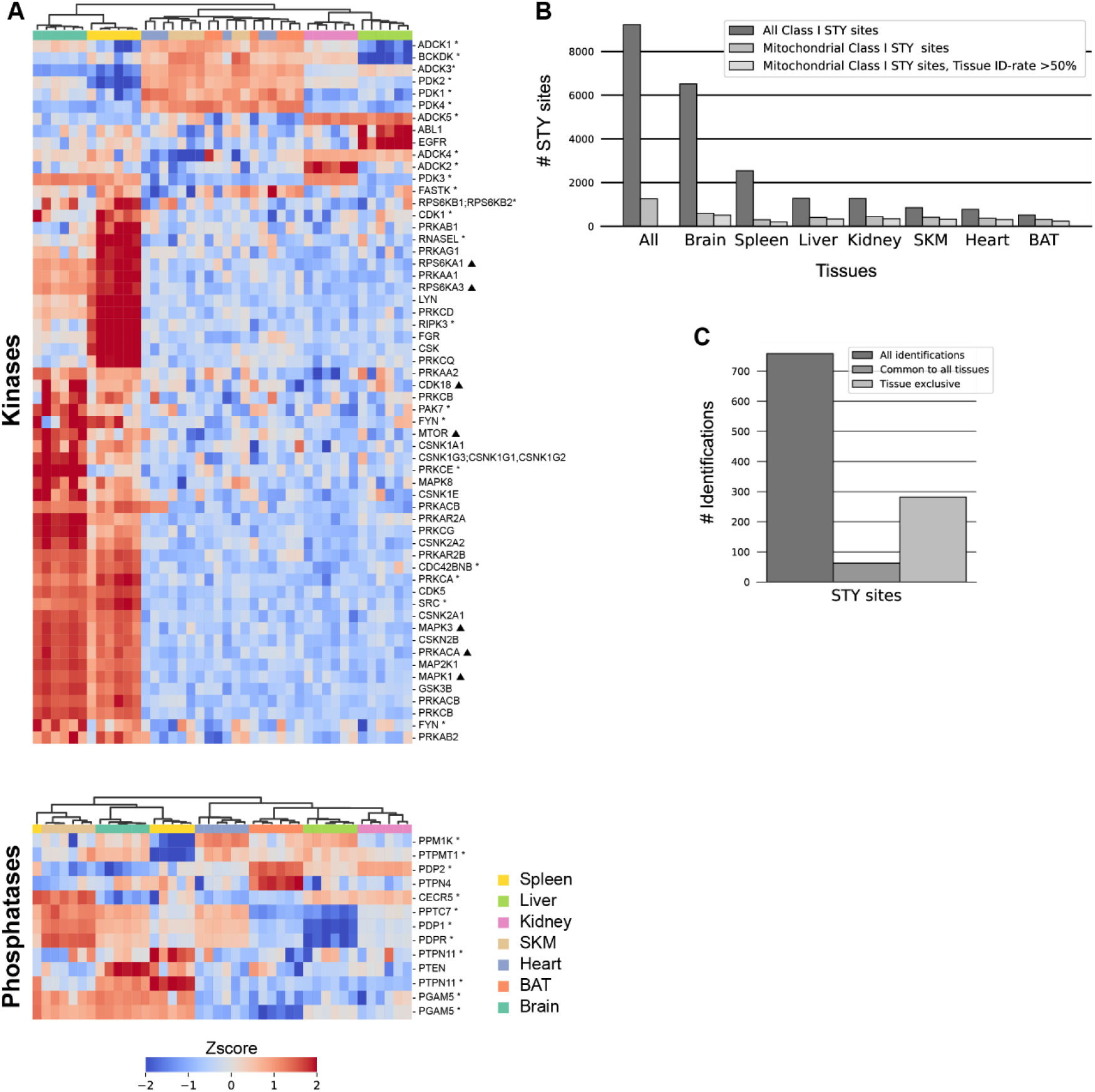
Tissue speceficity of mitochondrial kinases and phosphatases. (A) Z-scored protein abundances for predicted (triangle), known (star) (based on MitoCarta3.0 and IMPI database), and manually curated (Figure 5 - Source Data 1) mitochondrial kinases (top) and phosphatases (bottom) across analyzed tissues. Skeletal muscle (SKM), brown adipose tissue (BAT). (B) STY site identification numbers. (C) Mitochondrial STY site numbers after filtering for mitochondrial STY sites identified in at least 5 out of 6 biological replicates in one tissue. Identification numbers for all identified STY sites (left), STY sites common to all tissues (middle), and STY sites exclusive to one tissue (right) are shown. Data in this figure is based on the analysis of six replicates (n=6) for each tissue. Skeletal muscle (SKM), brown adipose tissue (BAT). See also Figure 5 – Source Data 1.

To investigate if and how kinase and phosphatase levels translate into protein phosphorylation, we analyzed the mitochondrial phosphoproteomes of the same samples collected from all seven tissues (Figure 1A). This analysis resulted in the identification of 1263 phosphorylation sites on 626 mitochondrial proteins (Figure 5B, Figure 5 - Source Data 1). After stringent filtering of the data for more than four identifications across six biological replicates in at least one tissue and a site localization score higher than 75%, we obtained a dataset of 758 phosphorylation sites on 423 mitochondrial proteins (Figure 5C, Figure 5 - Source Data 1). Of these high confidence sites, 16% have previously not been reported in mouse according to the PhosphoSitePlus (PSP) database (Hornbeck et al., 2012). Strikingly, in contrast to the mitochondrial proteomes, 37% of the phosphosites identified on mitochondrial proteins were exclusive to one tissue, and only 8% were identified in all the tissues measured (Figure 5C). This might indicate that mitochondrial diversity is more strongly pronounced at the phosphorylation than the protein level.

### Mitochondrial phosphoproteomes exhibit extensive intra-mitochondrial phosphorylation

To understand the distribution of mitochondrial phosphoproteins across mitochondrial compartments we examined the sub-mitochondrial localization based on the curated MitoCarta3.0 annotation. Mitochondria are typically divided into four main compartments, i.e. mitochondrial outer membrane (OMM), intermembrane space (IMS), inner mitochondrial membrane (IMM) and matrix, although the complex organization of the IMM possibly may define additional compartments (Colina-Tenorio et al., 2020). In line with the high proportion of shared mitochondrial proteomes across all tissues (Figure 1C), the overall localization of proteins was not different between tissues and closely resembled the distribution of all annotated mitochondrial proteins in the database (Figure 6A). However, when performing the same analysis using the phosphorylated mitochondrial proteins, we observed a significant shift towards a localization to the OMM in all tissues (adj. p-values <6.4 x 10^-10^)(Figure 6A, Figure 6 – supplement 1). Surprisingly, our data also showed that depending on the tissue type, more than 60% of phosphorylated mitochondrial proteins had an intra-mitochondrial annotation - IMM, IMS or matrix localization - (Figure 6A, Figure 6 – supplement 1). Interestingly, 36-54% of OMM, but only 7-22% of intra- mitochondrial proteins were phosphorylated. Here, especially brain (10%) and spleen (7%) tissues showed low intramitochondrial phosphorylation rates.

**Figure 6.**
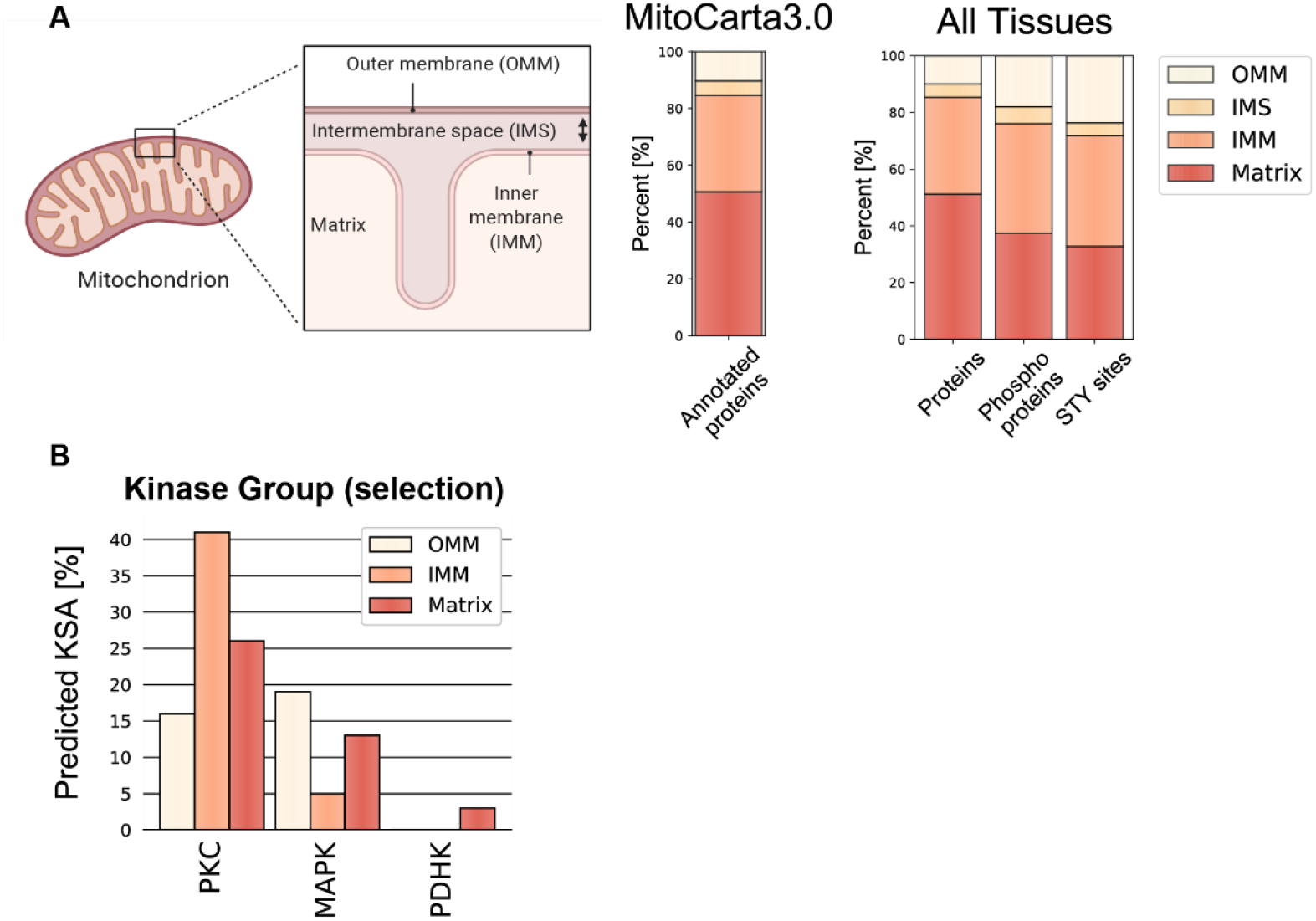
Localization distribution of mitochondrial phosphoproteome diverges from mitochondrial proteome. (A) Simplified scheme of a mitochondrion with four different mitochondrial localizations – outer mitochondrial membrane (OMM), intermembrane space (IMS), inner mitochondrial membrane (IMM), matrix – and the distribution of mitochondrial proteins contained in and classified by the MitoCarta3.0 database. Bar graphs show the precentral distribution of mitochondrial proteins (left), phosphoproteins (middle) and STY sites (right) across different mitochondrial localizations. (B) Predicted Kinase substrate associations (KSA) by the NetworKin3.0 tool for selected kinase families. Data in this figure is based on the analysis of six replicates for 7 different tissues. See also Figure 6 – Source Data 1.

This prompted us to further investigate the localization of specific kinase-substrate associations (KSA) across sub-mitochondrial localizations using NetworKin3.0 (Horn et al., 2014). Prominently, more than 40% of the predicted KSA in the IMM were linked to the PKC kinase family (Figure 6B). Studies have already reported the localization of PKC kinase family members to mitochondria, as well as an increased phosphorylation of the IMM protein COX IV after PKCɛ activation (Baines et al., 2002; Jaburek et al., 2006; Majumder et al., 2000; Ogbi and Johnson, 2006; Ping et al., 2002). Moreover, the MAPK group appeared to act on proteins localized to the OMM and matrix, while the PDHK family was specifically associated with the matrix proteins (Figure 6B). The members of the latter kinase family are known to localize to the mitochondria matrix (Hitosugi et al., 2011), further supporting the validity of identified KSA. However, molecular studies are needed to investigate such KSA, whether phosphorylation of intra-mitochondrial proteins occurs *in situ* or outside mitochondria before being imported into mitochondria, how and which kinases/phosphatases translocate to or into mitochondria and whether these phosphorylation events are functionally relevant.

### Mitochondrial phosphoproteome reveals tissue-specific modulation of fusion and fission events

The tissue-specific phosphorylation of mitochondrial proteins suggested functional differences in mitochondria and prompted us to investigate the influence of phosphorylation on mitochondrial dynamics. We focused on proteins involved in mitochondrial fusion and fission, two important counteracting events involved in organelle distribution, size balancing and maintenance of a healthy mitochondrial network (Liu et al., 2020; Silva Ramos et al., 2019). Especially proteins involved in the fission process are regulated by a range of protein modifications, including phosphorylation (Cribbs and Strack, 2007; Taguchi *et al*., 2007; van der Bliek et al., 2013).

Throughout all tissues, MIGA1 (FAM73A) and MIGA2 (FAM73B), two homologues regulating mitochondrial fusion by functioning downstream of the mitofusins, showed different abundances (Figure 6 – supplement 2). This is especially interesting since MIGA1 and MIGA2 can form hetero and homodimers, highlighting a different regulation of fusion in different tissues (Zhang et al., 2016). Moreover, we identified multiple phosphorylation site clusters on MIGA2, while none were identified on MIGA1 (Figure 6 – supplement 2). Intriguingly, similar phosphorylation clusters were identified on Miga in *Drosophila melanogaster* (Xu et al., 2020). Interestingly, two phosphorylation sites on Miga, S246 and S249, were reported to be essential for Vap33 interaction and the establishment of endoplasmic reticulum–mitochondria contact site (ERMCS), suggesting that phosphorylation on MIGA2 has similar functions in mammals (Xu *et al*., 2020).

Moreover, we observed that GTPase dynamin-related protein 1 (DRP1), a crucial player initiating mitochondrial fission (Bleazard et al., 1999; Cereghetti et al., 2008), displayed higher abundance in brain compared to other tissues (Figure 7A). This observation supports the importance of mitochondrial fission in neurons, where mitochondria switch to a fragmented morphology to enter and travel through axons (Lewis *et al*., 2018). Additionally, we also found elevated levels of serine 622 phosphorylation on DRP1 in brain tissue. This site has been shown to regulate DRP1 translocation to mitochondria (Cereghetti *et al*., 2008; Cribbs and Strack, 2007; Taguchi *et al*., 2007), indicating that it is actively localized to mitochondria, likely to regulate constant fission events in the brain tissue (Figure 7B).

**Figure 7.**
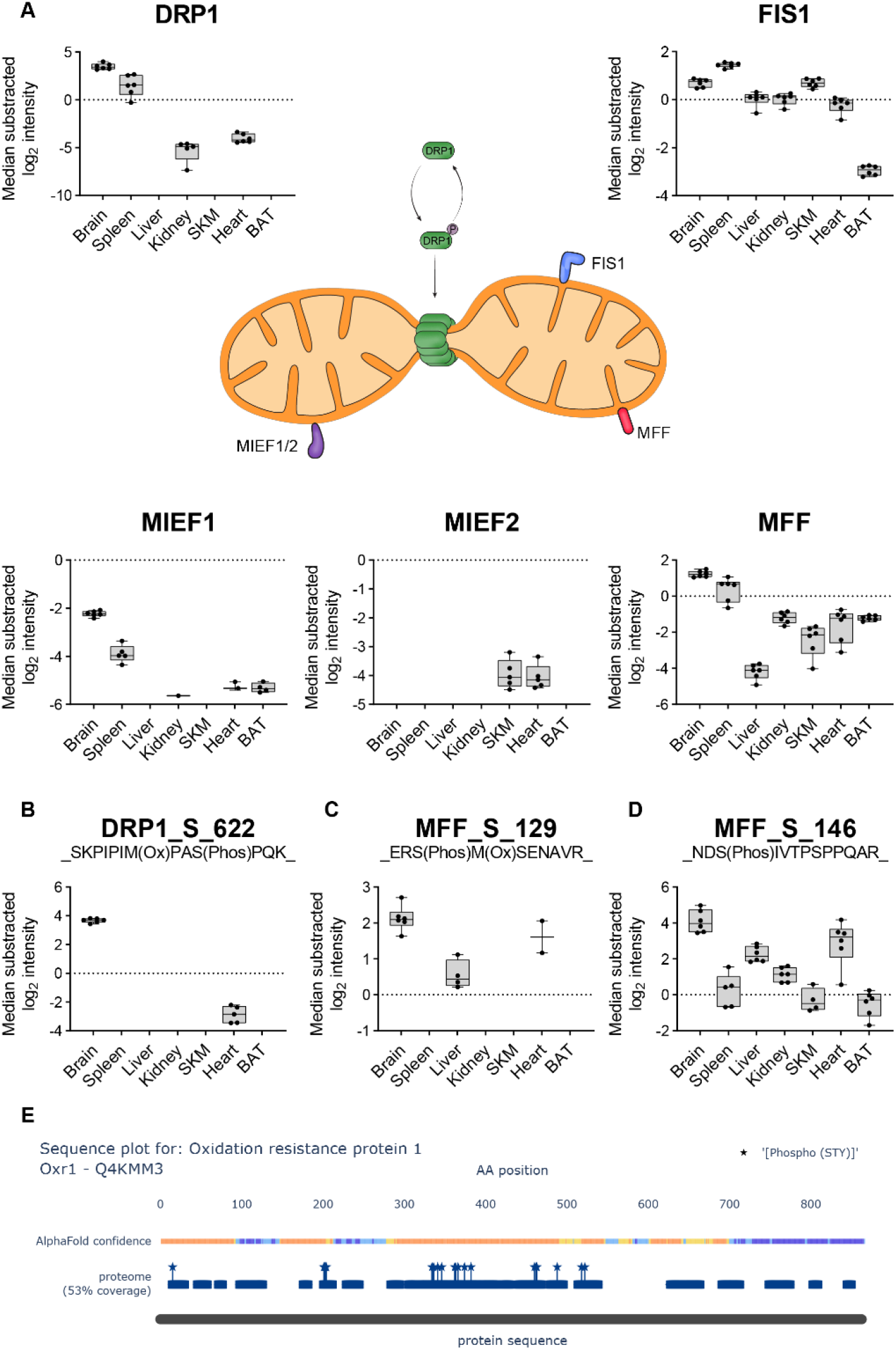
Phosphoproteome reveals tissue-specific functionality for mitochondrial fission. (A) Scheme displays the reversible phosphorylation and the connected localization change to mitochondria of DRP1 (green). Normalized intensities (median of all log2 transformed mitochondrial proteins of a sample was subtracted from all log2 protein intensities of that sample) of DRP1 across all analyzed tissues (black dots indicate individual identifications) are displayed in the upper left. Mitochondrial DRP1 receptors – FIS1 (blue), MIEF1/2 (violet), MFF (red) – and corresponding box plots are shown next to their receptors. (B) Normalized intensities (median of all log2 transformed mitochondrial phosphopeptide of a sample was subtracted from all log2 peptide intensities of that sample) of the phosphopeptide showing S622 phosphorylation on DRP1 (black dots indicate individual identifications). (C) Same as (B) showing S129 phosphorylation on MFF. (D) Same as (B) showing S146 phosphorylation on MFF. Significance of protein abundance differences in relation to the reference tissue (brain for DRP1 panel, brain and spleen in FIS1 and MFF panel) were estimated (p-values <0.001, except for SKM in FIS1 panel) by one-way ANOVA analysis (Figure 7 - Source Data 1). Significance of phosphosites abundance differences in (B) and (C, only brain and liver) were estimated by a two-side t-test (Figure 7 - Source Data 1). Significance of MFF_S_146 abundance differences in relation to the reference tissue (brain) were estimated (p-value<0.05) by one-way ANOVA analysis (Figure 7 - Source Data 1). Data in this figure is based on the analysis of six replicates (n=6). Skeletal muscle (SKM), brown adipose tissue (BAT). See also Figure 7 – Source Data 1.

In mammals, four DRP1 receptor proteins, that are all integral membrane proteins of the OMM, have been reported: mitochondrial fission protein 1 (FIS1), mitochondrial fission factor (MFF), and mitochondrial dynamics protein MiD49 (MIEF1) and MiD51 (MIEF2) (Loson et al., 2013). MFF and FIS1, the fission promoting receptors, were robustly quantified in all tissues and displayed higher abundances in brain and spleen compared to other tissues (Figure 7A). However, MIEF1/2, which counteract DRP1- mediated fission (Dikov and Reichert, 2011; Liu et al., 2013), were generally too low to be robustly quantified in the measured tissues. Additionally, we detected higher levels of MFF phosphorylation at the serine 129 and 146 residues in brain tissue compared to all other tissues in which they were detected (Figure 7C and 7D). Both sites are essential for the recruitment of DRP1 and initiation of fission (Ducommun *et al*., 2015; Lewis *et al*., 2018; Toyama *et al*., 2016).

Elevated DRP1 levels have been shown to increase ROS levels (Watanabe et al., 2014). Intriguingly, we identified oxidation resistance 1 (OXR1), exclusively in the mitochondria of brain tissue, where it plays an important role in the protection of neuronal cells from oxidative stress (Volkert and Crowley, 2020). This likely indicates a protective function of OXR1 in brain tissue as a response to the prevalent fragmented organellar morphology induced by DRP1. In addition, we found OXR1 to be hyperphosphorylated and three out of 12 high confidence sites were novel (Figure 7E). Tissue specificity and the lack of functional annotation of phosphorylation sites that are identified in our study make OXR1 an exciting candidate to be investigated in the future.

### Web application makes Mouse Mitochondria Atlas data readily accessible

As indicated by the above examples, this study presents a rich resource to explore the mitochondrial proteomes and phosphoproteomes across mouse tissues. Preceding examples show the potential of this resource for investigation of tissue-specific mitochondrial regulations on the proteome and phosphoproteome level, ultimately permitting the generation and analysis of new hypotheses. However, utilization of such resources largely depends on the ease of data access for exploration.

To this end, we created a web application mitophos.de offering the end user an interface to easily explore datasets, including MitoCarta3.0 networks, abundance comparisons across tissues and sequence analysis (Figure 8A). As an example, Figure 8 illustrates the MICOS complex, which has a central role in mitochondria (Khosravi and Harner, 2020). In the network view one can see (I) members of the complex as well as the phosphorylation sites identified on these proteins and (II) in which tissues and how reproducibly they are identified in our dataset (Figure 8B). Moreover, the user can inspect individual abundance distributions of all identified proteins/STY sites across all measured tissues. For instance, MINOS1, a core component of the MICOS complex (Bohnert et al., 2015), displayed high abundance in mitochondria isolated from heart, SKM and BAT tissues (Figure 8C). Moreover, in the sequence view the AlphaMap tool (Voytik et al., 2021) is integrated to map all identified peptides, including phosphorylated peptides, onto their respective protein sequence along with structural information such as topological domains and transmembrane regions (Figure 8D) and to visualize phosphorylation sites in their 3D structures (unpublished data) as predicted by AlphaFold (Jumper et al., 2021).

**Figure 8.**
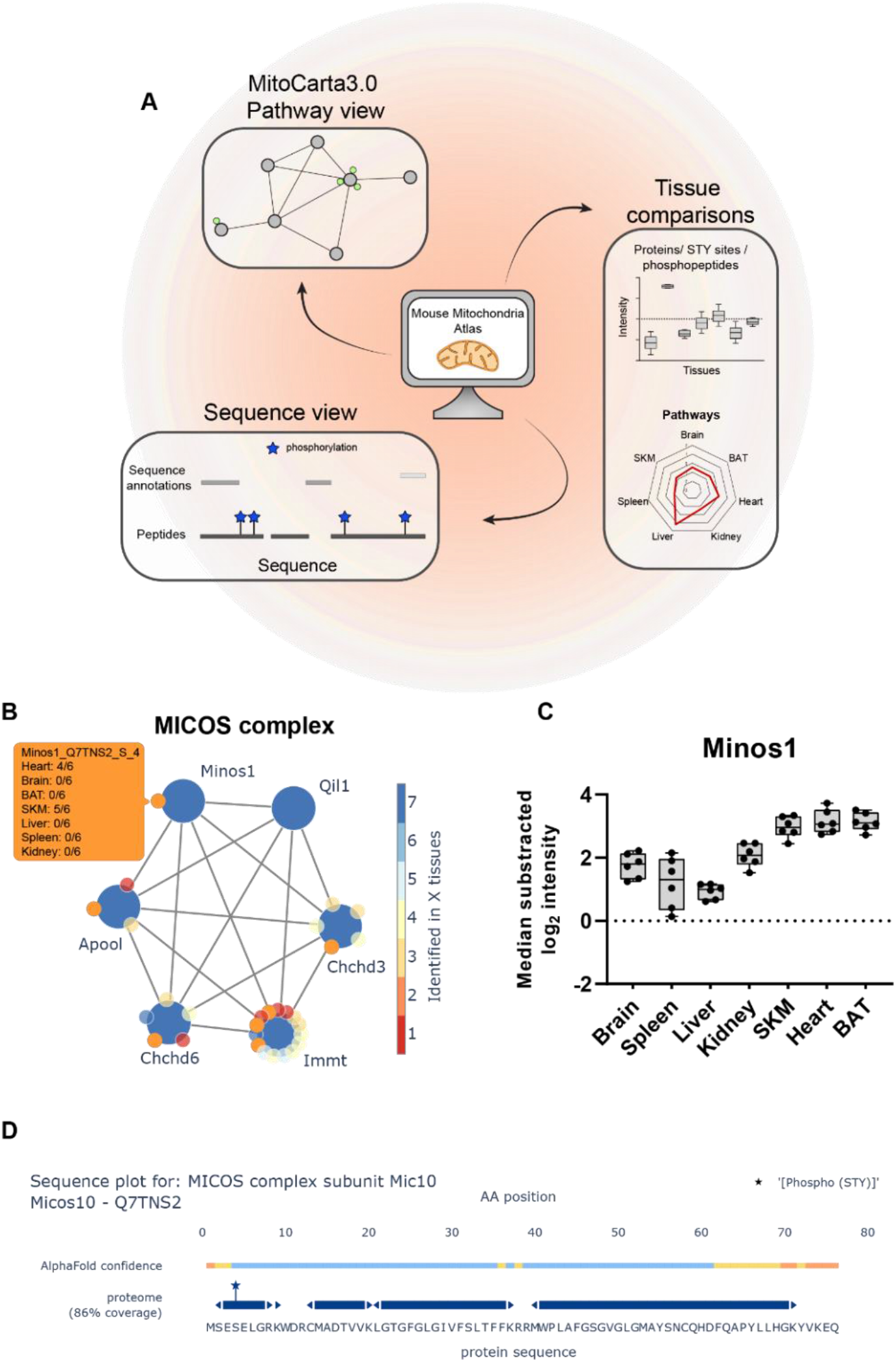
Web application readily enables easy data access. (A) Scheme of Mouse Mitochondria Atlas application features (<u>mitophos.de</u>). (B) MICOS complex view based on MitoCarta3.0 annotation. Large and small nodes represent proteins and class I STY sites, respectively. Edges represent on String interaction scores >0.4 and color of nodes indicate the number of tissues in which proteins/STY sites were identified. (C) Normalized intensities (median of all log2 transformed mitochondrial proteins of a sample was subtracted from all log2 protein intensities of that sample) for Minos1 across all analyzed tissues (black dots indicate individual identifications, box and error bar). (D) Sequence plot of Micos1 shows structural information (based on UniProt annotations) and protein coverage based on identified peptides. All identified STY sites are marked with a star. Data in this figure is based on the analysis of six replicates (n=6). Skeletal muscle (SKM), brown adipose tissue (BAT).

Together, this data-rich and comprehensive tool is an entry point to investigate the herein presented resource and will assist in future efforts to functionally characterize mitochondrial proteins and their respective phosphorylation sites.

## Discussion

Here we present a tissue-specific atlas of mouse mitochondrial proteomes and phosphoproteomes, an in-depth resource towards a better understanding of the composition and function of this vital organelle in a tissue-specific manner. Previous MS-based studies combining mitochondrial phosphoproteome and proteome measurements typically focused on a single or few tissues and were generally shallower than our study. In addition, differences in study designs such as utilization of various organisms or mitochondria/phosphopeptide enrichment protocols, analysis pipelines, and mitochondrial protein annotation databases, complicate the integration of such datasets to understand tissue specificity. We now globally and precisely quantified different protein expression and phosphorylation patterns at subcellular level across seven mouse tissues, providing a detailed view on mitochondrial diversity. The breadth and depth of coverage achieved by the integration of tissue-specific mitochondrial proteomes and phosphoproteomes provide unbiased insights into the mitochondrial composition and function. This allows the generation and assessment of novel hypotheses related to mitochondrial biology, which cannot be generated with focused studies alone. Integrated mitochondrial proteomes and phosphoproteomes that are diverse between tissues can readily be explored at mitophos.de.

Our data revealed that the functional diversity of mitochondria is defined by protein abundance rather than compositional differences since more than half of the mitochondrial proteome was shared by all analyzed tissues and 90% by at least two tissues. For instance, the electron transport chain is an integral part of the mitochondrial composition and its components are found across all tissues, however, their abundance shows substantial differences to meet tissue-specific energy demands. Thus, dysregulation of individual proteins can strongly affect mitochondria in one tissue, leading to severe diseases, while mitochondria in a different tissue remain largely unaffected. Moreover, we found that 9% of the proteome displays tissue specificity, further contributing to our understanding of tissue-specific effects of mitochondrial protein dysregulation (Russell et al., 2020). This is an important concept, which can now be studied in an unbiased manner using our resource data, providing opportunities for development of targeted treatments for mitochondrial diseases. For instance, members of the SLC25 family are linked to metabolic diseases in distinct tissues (Palmieri and Monne, 2016) and various cancers (Rochette et al., 2020). However, biological functions of a large repertoire of mitochondrial SLCs are still unknown. For example, inactivation of SLC25A25 was assessed in mouse skeletal muscle tissue where it caused a reduced metabolic efficiency (Anunciado-Koza et al., 2011). Given its high abundance in mitochondria of brain tissue and in glioma cells (Traba et al., 2012), it will also be interesting to investigate its role in this tissue, particularly whether SLC25A25 deficiency influences neuronal fitness.

Mitochondria are essential cellular entities that are involved in a wide variety of cellular processes (McBride et al., 2006) through dynamic interaction and constant communication with other organelles such as the (ER), nucleus and peroxisomes via membrane contact sites (Desai et al., 2020; Perrone et al., 2020; Shai et al., 2018) or protein complexes, such as the ribosome (Lashkevich and Dmitriev, 2021). We suggest that non-mitochondrial proteins identified in the samples might present signatures that could convey important biological information regarding mitochondria-associated structures. Given the high level of mitochondrial enrichment combined with highly reproducible LC-MS/MS measurements, such protein signatures are unlikely to be solely based on unspecific enrichment of abundant proteins. For example, the proteasome is robustly identified in most of the mitochondria enriched samples, which is in line with its involvement in the degradation of misfolded mitochondrial proteins (Basch et al., 2020; Kodron et al., 2021). We also prominently observed that several members of the cytochrome P450 superfamily are enriched in liver mitochondria, demonstrating the versatile interaction of mitochondria with their cellular environment. Furthermore, our data identified non-mitochondrial ribosomal proteins in all tissues, which could be explained by the local translation of nuclear-encoded mitochondrial mRNAs (Lashkevich and Dmitriev, 2021). It was recently shown that RNA-bearing late endosomes associate with mitochondria and ribosomes forming hotspots of local protein synthesis in axons (Cioni et al., 2019) and that mitochondria fuel such local translation in neurons, enabling synaptic plasticity (Rangaraju et al., 2019).

There is mounting evidence that phosphorylation of mitochondrial proteins fulfills important functions to maintain cellular health as exemplified by the fission process in this study. Deregulation of mitochondrial protein phosphorylation can lead to diseases such as cancer, diabetes, heart and neurological disorders. It was recently reported that 91% of mitochondrial proteins on MitoCarta3.0 have at least one phosphorylation site reported on the PSP database (Niemi and Pagliarini, 2021). However, this analysis also includes proteins that do not always localize to mitochondria (Ben-Menachem et al., 2011) and phosphorylation sites that can specifically be captured upon perturbations and stimuli. Our study revealed that around half of the mitochondria-localized proteins were phosphorylated in tissues at steady state - 39% in brain, 15% in spleen, 29% in liver, 28% in kidney, 32% in SKM, 31% in heart and 22% in BAT. This suggests that the mitochondrial proteome and phosphoproteome compositions are dynamically modulated in response to environmental changes. Furthermore, we identified over 60 kinases and 10 phosphatases that either are localized to mitochondria or associate with mitochondria, providing a global view on important modulators of mitochondrial protein phosphorylation. Mapping tissue-specific mitochondrial kinases and phosphatases is an important step towards understanding their role in the regulation of the mitochondrial phosphoproteome in different tissues and hence developing therapeutics for mitochondrial diseases. For example, a mitochondrial phosphatase, phosphoglycerate mutase family member 5 (PGAM5), has recently emerged as an important regulator of mitochondrial homeostasis. Deletion of PGAM5 has been shown to result in Parkinson’s-like movement disorder in mice (Lu et al., 2014) and T cell dysfunction in primary cells (Panda et al., 2016). While its diverse roles largely remain to be uncovered (Liang et al., 2021), a novel PGAM5 inhibitor was recently suggested as a potential therapeutic for brain ischemic stroke (Gao et al., 2021). We observed that PGAM5 displays elevated levels in the mitochondria isolated from brain, SKM and spleen, possibly explaining the tissue-specific phenotypes induced by its absence and where in the body molecules targeting its phosphatase activity would exert their effects.

The data in this study will contribute to our understanding of the tissue-specific composition and function of mitochondria and serve as a gateway for investigation of specific questions related to mitochondrial biology. Future biochemical and more focused investigations are needed to validate our findings and test the hypotheses arising from our study. For instance, it is pivotal to experimentally validate kinase- substrate associations of previously unknown phosphorylations on mitochondrial proteins and their functional implications in the cell. Additonally, the impact of those phosphorylations on the localization of target proteins and, more specifically, the question of wheter mitochondrial proteins are phosphorylated before or after entering the mitochondria remain to be investigated. Accessibility, for instance, of previously undescribed phosphorylation sites on mitochondrial proteins can be assessed using advanced structural tools (Jumper *et al*., 2021) to determine if they are likely targeted by a mitochondrial kinase or a cytoplasmic kinase before being imported (Schober et al., 2021). Moreover, future developments towards better enrichment strategies for the isolation of mitochondria from different tissues and advances in the MS technology will aid to further improve the depth and quality of the mitochondrial proteomes and phosphoproteome.

## Materials and Methods

### Experimental model and subject details

Six C57BL/6N mice (3male, 3 female) were housed in a 12-hours light/dark cycle in standard ventilated cages under specific-pathogen-free conditions with constant temperature (21°C) and humidity (50 - 60%) and fed ad libitum with a standard mouse diet. At the age of 18-21 weeks mice were sacrificed by cervical dislocation. The study was approved by the by the Landesamt für Natur, Umwelt und Verbraucherschutz Nordrhein–Westfalen, Germany, and performed in accordance with European law.

### Tissue preparation and isolation of ultra-pure mitochondria

Mice were sacrificed by cervical dislocation and the 7 tissues - heart, skeletal muscle (SKM), brown adipose tissue (BAT), spleen, kidney, liver, and brain – were rapidly removed. Heart, spleen, and kidney, tissues were homogenized in mitochondrial isolation buffer containing 320 mM sucrose, 1 mM EDTA, and 10 mM Tris-HCl, pH 7.4, supplemented with EDTA-free complete protease inhibitor cocktail and PhosSTOP tablets (Roche). For isolation of mitochondria from BAT, liver, and brain the mitochondrial isolation buffer was additionally supplemented with 0.2% bovine serum albumin (Sigma-Aldrich). Subsequently, crude mitochondria were isolated from the homogenates by two rounds of differential centrifugation (see Figure 1a). Isolation of crude mitochondria from SKM was performed as previously described (Frezza et al., 2007)). Crude mitochondrial pellets from all tissues were further purified on a Percoll density gradient as described recently (Kuhl *et al*., 2017). Briefly, mitochondrial pellets were washed once in 1xM buffer (220 mM mannitol, 70mM sucrose, 5mM HEPES pH 7.4, 1 mM EGTA pH 7.4; pH was adjusted with potassium hydroxide; supplemented with EDTA-free complete protease inhibitor cocktail and PhosSTOP tablets (Roche)) and subsequently purified on a Percoll (GE healthcare) density gradient of 12%:19%:40% via centrifugation in a SW41 rotor at 42, 000 g at 4°C for 30 min in a Beckman Coulter Optima L- 100 XP ultracentrifuge using 14 mm × 89 mm Ultra-Clear Centrifuge Tubes (Beckman Instruments Inc.). Ultra-pure mitochondria were harvested at the interphase between 19% and 40% and washed three times with 1xM buffer. Dry mitochondrial pellets were snap-frozen in liquid nitrogen and stored at -80°C until further use.

### Mitochondrial (phospho)proteome sample preparation

Frozen ultra-pure mitochondria pellets were resuspended in lysis buffer (4%SDC, 100mM Tris/HCl, pH8.5), boiled for 5 min at 95°C and sonicated in 30 s intervals for 15 min (Bioruptor). Protein concentration was estimated via Tryptophan assay (Kulak et al., 2014) and was adjusted with lysis buffer to a total volume of 270 µl containing 400 µg of protein for brain, SKM, liver, heart, kidney, and 140 µg of protein for BAT samples. Proteins were reduced and alkylated by adding 30 µl of 10x reduction/alkylation solution (100 mM Tris (2-carboxyethyl)phosphine hydrochloride (TCEP) and 400 mM 2-chloroacetamide (CAA), followed by 5min incubation at 45°C. Subsequently, 1:100 Trypsin and LysC were added for overnight protein digestion at 37°C. For proteome analysis 10 µl (brain, SKM, liver, heart, kidney) and 20 µl (BAT, spleen) aliquots were taken and loaded on SDB-RPS StageTips. Peptides were washed with 200µl wash buffer (0.2% TFA/2% ACN (vol/vol)) and then eluted with SDB-RPS elution buffer (1.25% NH_4_OH, 80% ACN (vol/vol)) and dried in a SpeedVac. Dried peptides were resuspended in A* buffer (2% ACN/0.1% TFA).

The remaining samples were processed following the EasyPhos protocol for phosphopeptide enrichment (Humphrey *et al*., 2018). In brief, samples were first mixed with isopropanol and EP buffer (48% TFA, 8 mM KH_2_PO_4_), followed by phosphopeptide enrichment with 5mg TiO_2_ beads per sample (GL Sciences). For this, samples were mixed with TiO_2_ beads in loading buffer (6% TFA/80% ACN (vol/vol)) at a concentration of 1 mg/µl and incubated for 5 min at 40°C by shaking at 1200rpm. Subsequently beads were washed four times with 1 ml of wash buffer (5% TFA,60% isopropanol (vol/vol)) and phosphopeptides were eluted from beads using 60 µl of elution buffer (40% ACN, 5% NH_4_OH) and concentrated in a SpeedVac for 30 min at 45°C. Samples were immediately diluted with 100 µl of SDBRPS loading buffer (99% isopropanol, 1% TFA (vol/vol)) and loaded on SDB-RPS StageTips. Thereafter, phosphopeptides were washed and eluted as described above and resuspended in 6 µl A*.

### LC-MS/MS

For all measurements peptides were loaded onto a 50cm, in-house packed, reversed-phase column (75µm inner diameter, 1. diameter, ReproSil-Pur C18-AQ 1.9 μm resin [Dr. Maisch GmbH]) and separated with and binary buffer system consisting of buffer A (0.1% formic acid (FA)) and buffer B (0.1% FA in 80% ACN). The column temperature was controlled by a homemade column oven and maintained at 60°C. For nanoflow liquid chromatography an EASY-nLC 1200 system (Thermo Fisher Scientific), directly coupled online with a Q Exactive HF-X (Thermo Fisher Scientific) via a nano-electrospray source, was operated at a flow rate of 300 nl/min and 350 nl/min for mitochondrial proteome and phosphoproteome measurements, respectively.

For mitochondrial proteome measurements 500µg of peptides were loaded and separated using a gradient starting at 5% buffer B, increasing to 30% buffer B in 80 min, 60% buffer B in 4 min and 95% buffer B in 4 min. The MS was operated in DDA mode (Top12) with a full scan range of 300-1650 m/z and a MS1 and MS2 resolution of 60,000 and 15,000, respectively. The automatic gain control (AGC) was set to 3e6 and 1e5 for MS1 and MS2, while the maximum injection time was set to 20 ms and 60 ms, respectively. Precursor ion selection width was kept at 1.4 m/z and fragmentation was achieved by higher-energy collisional dissociation (HCD) (NCE 27%). Dynamic exclusion was enabled and set to 20 s.

For mitochondrial phosphoproteome measurements, 5 µl as loaded and separated using a gradient starting at 3% buffer B, increasing to 19% buffer B in 40 min, 41% buffer B in 20 min and 90% buffer B in 5 min. The MS was operated in DDA mode (Top10) with a full scan range of 300-1600 and a MS1 and MS2 resolution of 60,000 and 15,000, respectively. The automatic gain control (AGC) was set to 3e6 and 1e5 for MS1 and MS2, while the maximum injection time was set to 120 ms and 60 ms, respectively. Precursor ion selection width was kept at 1.6 m/z and fragmentation was achieved by higher-energy collisional dissociation (HCD) (NCE 27%). Dynamic exclusion was enabled and set to 30 s.

### Raw data analysis

DDA raw data were analyzed with MaxQuant (1.6.14.0) against the mouse fasta file (downloaded 19. October 2020) using default settings. PSM and protein dales discovery rate were controlled at 1% FDR. The match between runs (MBR) functionality was enabled and set in a way that only biological replicates belonging to the same tissue type were allowed to match each other. This eliminates the possibility of potentially false MBR identification transfer between tissues. Carbamidomethyl (C) was selected as fixed modification and Acetyl (Protein N-term) and oxidation (M) were defined as variable modifications. For mitochondrial phosphoproteome analysis, STY site phosphorylation was additionally selected as variable modification.

### Bioinformatics analysis

Data analysis was performed using the python programing language using python (3.8.12) and the following packages: alphamap, matplotlib (3.5.0), mygene (3.2.2), numpy (1.19.2), pandas (1.1.3), pyteomics (4.3.3), requests (2.26.0), scipy (1.7.2), seaborn (0.11.2), sklearn (0.0), upsetplot (0.6.0). All notebooks used for data analysis are available at GitHub (https://github.com/MannLabs). Identified proteins were filtered for at least 3 valid values in at least one tissue. Similarly, phosphorylation sites were filtered for at least 5 valid values in at least one tissue and a localization probability >75%. The UniProt API was used to map ‘ACC+ID’ protein group identifiers provided by the MQ analysis to ‘ENSEMBL_ID’, ‘P_ENTREZGENEID’ and ‘STRING_ID’ for further analysis. ‘ENSEMBL_ID’ and ‘P_ENTREZGENEID’ identifier were used for mitochondrial protein annotation based on the IMPI (IMPI_2020_Q3, downloaded 27. October 2020) and MitoCarta3.0 (downloaded 1. January 2021) databases, respectively. Network analysis and visualization were performed with the StringApp (1.6.0) in Cytoscape (3.8.2). Kinase annotations are based on manual annotations (Figure 6 - Source Data 1) and ‘pkinfam’ (downloaded 9. Arpil 2021). Networkin3.0 was used for kinase substrate association (KSA) predictions, while the Networkin score was set to 1. Gene Ontology (GO) annotations for the GOBP term enrichment analysis were retrieved from UniProt (accessed 8. March 2021). Enrichment analysis was performed in Perseus (1.6.7.0) against the set of identified proteins in the corresponding tissue and the results were filtered for an intersection size >10. Missing values were only imputed for PCA and heatmap analysis. For this, a Gaussian normal distribution with a width of 0.3 relative to the standard deviation of measured values and a downshift of 1.8 standard deviations were used. For data normalization, intensity values were log_2_ transformed and then filtered for known and predicted mitochondrial proteins. The median value of these mitochondrial proteins was subtracted from all log_2_ transformed values. Significance testing for individual proteins and phosphopeptides as shown in Figure 3 and Figure 7 was performed with the ordinary one-way ANOVA method or by two-sided t-tests in GraphPad Prism (9.3.1) (Figure 3 - Source Data 1, Figure 7 - Source Data 1). Significance testing for differences in mitochondrial protein, phosphoprotein and phosphosite localization was performed in RStudio (1.3.1093) (Figure 6 - Source Data 1). OMM proportions were used for fitting a beta-regression model using the betareg R package with default settings (Cribari-Neto and Zeileis, 2010; Grun et al., 2012). P-values were estimated with the lrtest function of the lmtest package (Zeileis and Hothorn, 2015) and p-values were adjusted with the “fdr” method of the p.adjust function of the stats base package (Benjamini and Hochberg, 1995)

### Website tool

The website tool is structured into four sections. The first three ‘Pathway view’, ‘Sequence view’, and ‘Tissue comparison’ are for displaying data, while the fourth section provides explanations for each individual section. In the ‘Pathway view’ and ‘Tissue comparison’, proteome and phosphoproteome data filtered for at least three and 5 identifications in at least one tissue, respectively. Intensity values were normalized as described above and used for data representation in the ‘Tissue comparison’ tab or z-scored across tissues and used for the ‘Pathway view tab’. Here, median z-score values of the six biological replicates per tissue are displayed in the data table. The polar plot represents the median z-score of all pathway/complex members of a given tissue. Network/complex annotations were retrieved from MitoCarta3.0 and protein interactions are based on STRING interaction scores. These STRING interaction scores were retrieved from STRING (17. November 2021) using the ‘STRING_ID’ and the STRING API. For the ‘Sequence view’, the ‘evidence.txt’ of the MaxQuant output files was directly used as input to annotated sequences.

The python programming language was used for data processing and visualization for the Dashboard. The following libraries were used for data processing: numpy (1.19.2), pandas (1.19.2), re, sys, os, and pyteomics (4.3.3). Several libraries from the HoloViz family of tools were used for data visualization and creation of the dashboard, including panel (1.14.6), holoviews (1.14.6), bokeh (2.2.2), plotly (4.12.0), and param (1.10.0). Network visualization was achieved with the NetworkX package (Hagberg et al., 2008). The Alphamap tool (Voytik *et al*., 2021) was integrated to display linear protein sequence annotations as well as to visualize 3D protein structures.

### Resource availability

#### Data and code availability

Datasets generated in this study have been deposited at ProteomeXchange and are publicly available as of the date of publication. The accession number is listed in the key resource table. (Identifier: PXD030062).

All original code has been deposited on GitHub (https://github.com/MannLabs) and is available as of the date of publication.

Any additional information required to reanalyze the data reported in this paper is available from the lead contact upon request.

## Acknowledgments

This work was supported by grants to N.G.L from the Swedish Research Council (2015-00418), Swedish Cancer Foundation (2021.1409), the Knut and Alice Wallenberg foundation, European Research Council (ERC Advanced Grant 2016-741366), grants from the Swedish state under the agreement between the Swedish government and the county councils (SLL2018.0471). L.S.K was supported by an EMBO long-term fellowship (ALTF 570-2019). I.K. was supported by the French Muscular Dystrophy Association (AFM-Téléthon #23294) and Agence nationale de la recherche (ANR-20-CE12-0011). We thank all members of the Department of Signal Transduction and Proteomics, in particular, Isabel Bludau for helpful discussions regarding the website tool, Igor Paron for MS technical assistance, Mario Oroshi for technical assistance during website setup, Marta Murgia for helpful discussions on mitochondrial biology and Johannes Müller-Reif, Patricia Skowronek and Sophia Steigerwald for columns. We are grateful to Jesper Olsen and Brenda Schulman for constructive and insightful discussions. Figure 1A and Figure 6A were partly created with BioRender.com.

## Author contributions

F.M.H., L.S.K. designed experiments. Mouse work was performed by L.S.K. and I.K.. Proteomic experiments were conducted by F.M.H. and O.K.. Website was constructed by F.M.H. and I.B.. Data were analyzed by F.M.H.. All authors contributed to writing and editing of the manuscript.

## List of Source Data

Figure 1 – Source Data 1

Figure 1 – Source Data 2

Figure 1 – Source Data 3

Figure 1 – Source Data 4

Figure 1 – Source Data 5

Figure 1 – Source Data 6

Figure 1 – Source Data 7

## List of Supplementary Files

Supplementary File 1 Coefficient of variation and dynamic range of protein identification

Supplementary File 2 Mitochondrial protein intensity ranking

## Supplementary Figures

**Figure 2 – figure supplement 1.**
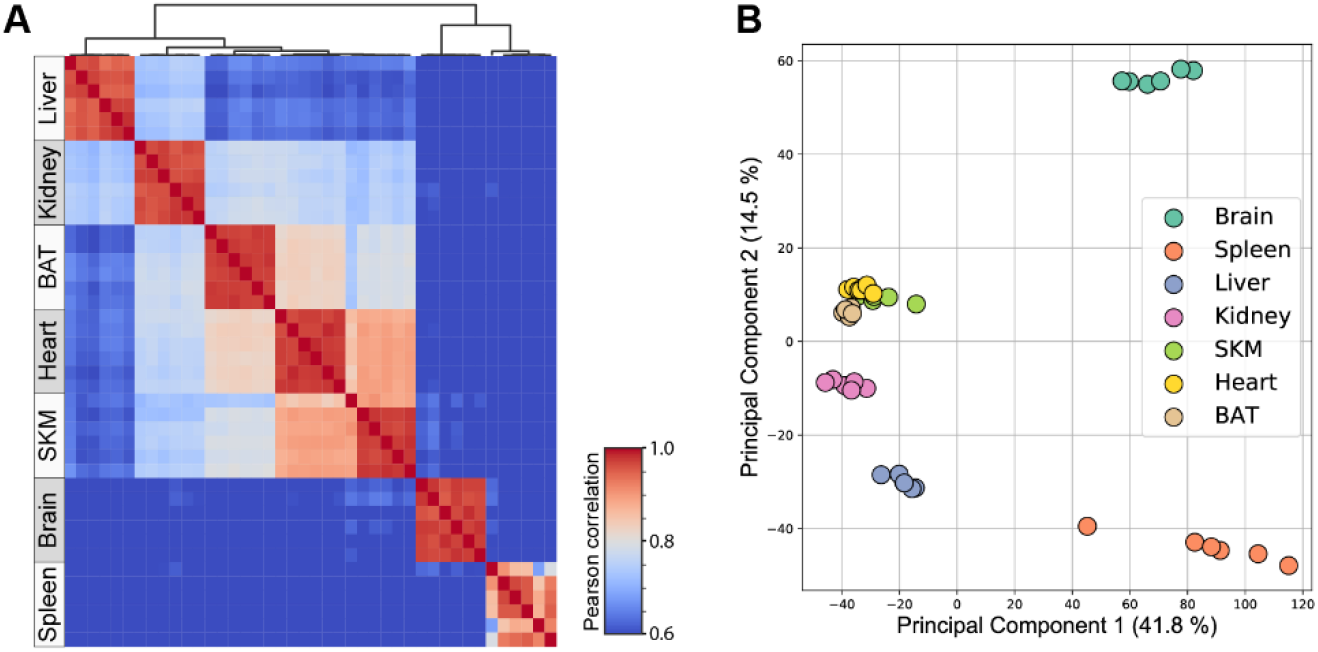
Mitochondria enriched samples show tissue-specific clustering. (A) Heatmap showing Pearson correlations for biological replicates (n=6) for proteins of all mitochondrial enriched samples. (B) Principal component analysis of all proteins of all acquired biological replicates (n=6) (Figure 2 – Source Data 1). Skeletal muscle (SKM), brown adipose tissue (BAT).

**Figure 4 – figure supplement 1.**
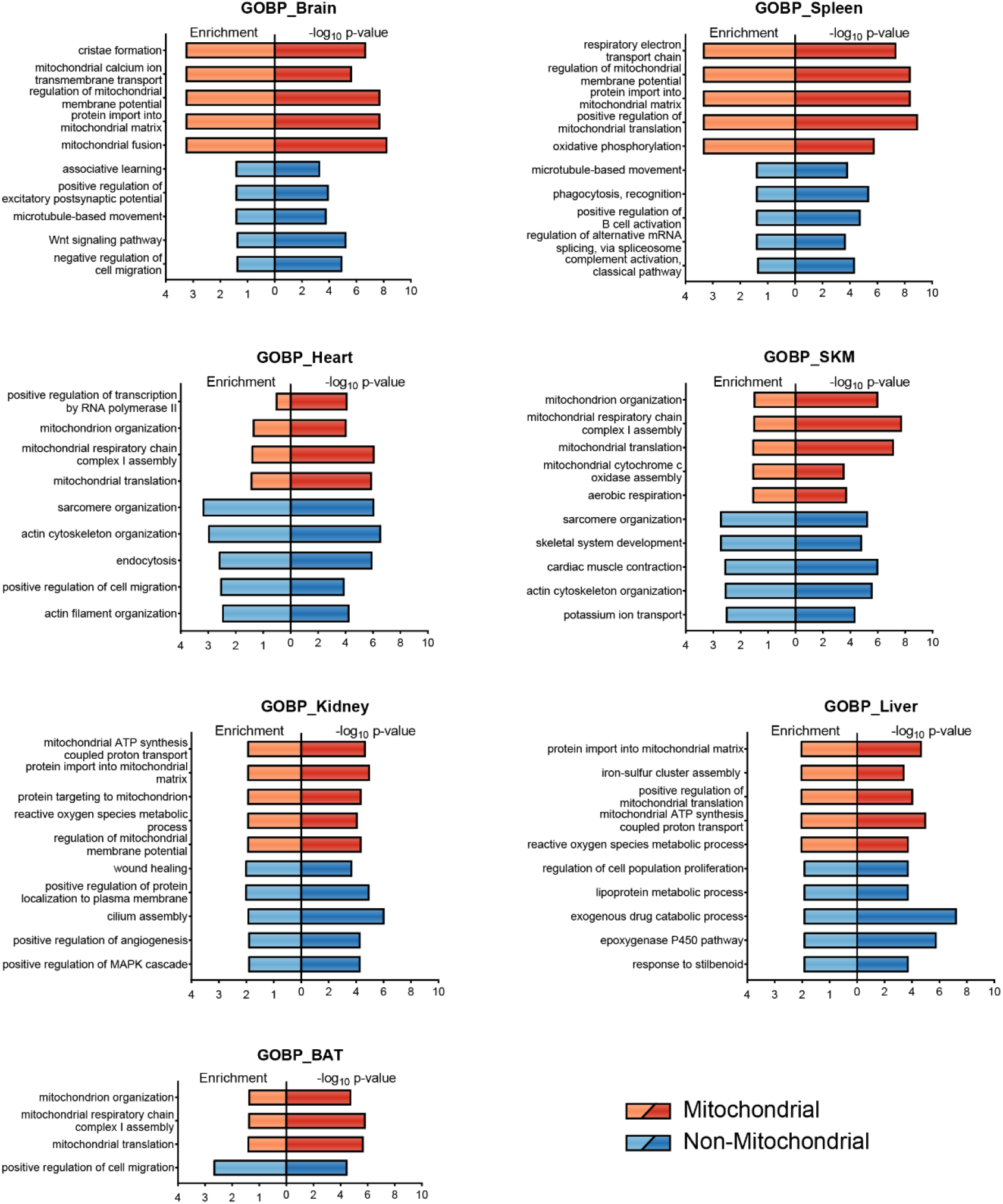
Gene Ontology enrichment displays tissue specificity. Geno Ontology biological process (GOBP) enrichment for mitochondrial (orange) and non-mitochondrial (blue) proteins. Enrichment analysis was performed in Perseus (1.6.7.0) against the set of identified proteins in the corresponding tissue and the results were filtered for an intersection size >10 and the top 5 enriched terms of each tissue are displayed (Figure 4 – Source Data 1). Skeletal muscle (SKM), brown adipose tissue (BAT).

**Figure 6 – figure supplement 1.**
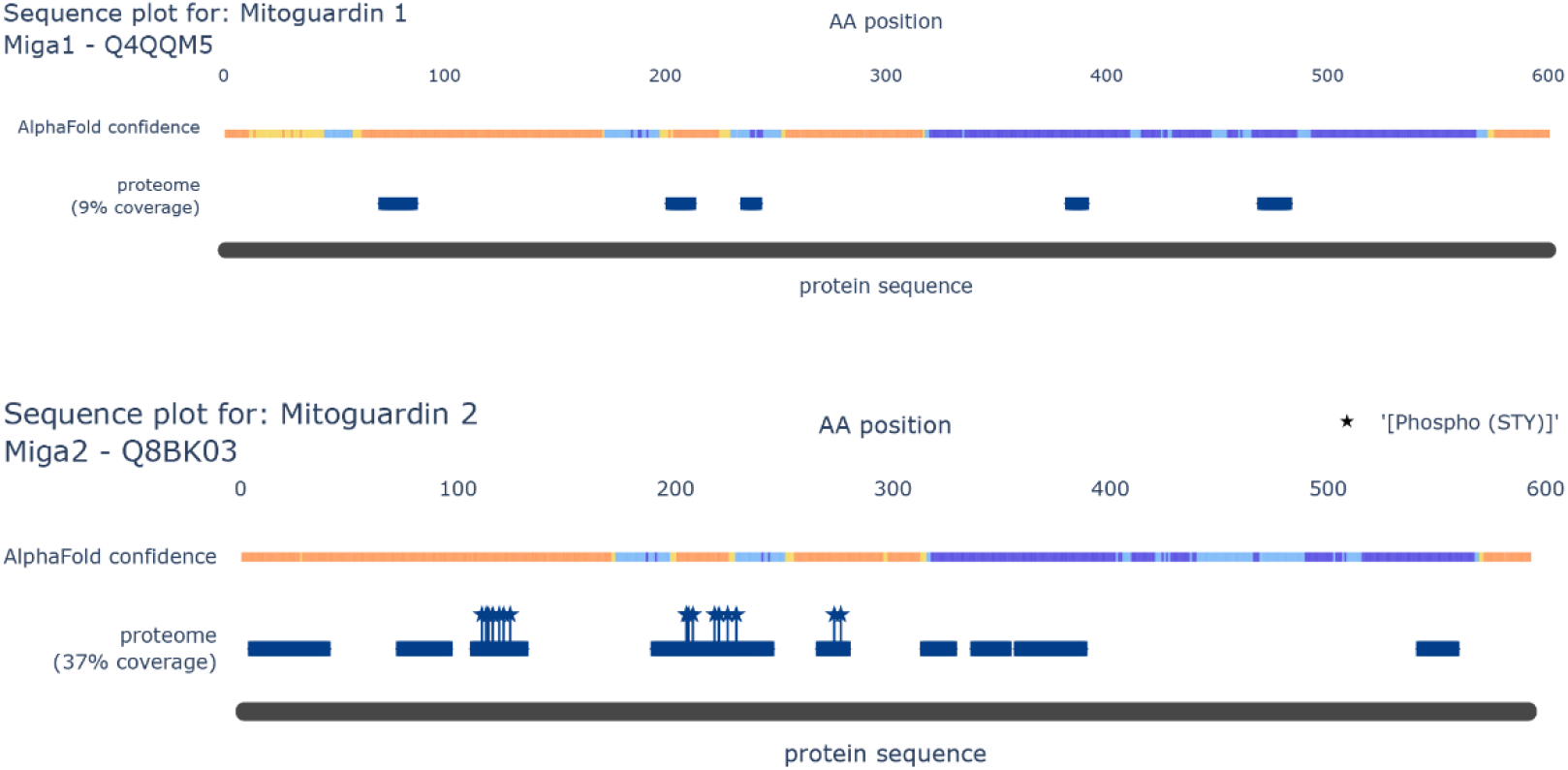
Sequence annotation highlights phosphorylation clusters on MIGA2. Sequence plot of MIGA1 (top panel) and MIGA2 (bottom panel) show structural information (AlphaFold confidence scores) and protein coverage based on identified peptides. All identified STY sites are marked with a star.

**Figure 6 – figure supplement 2.**
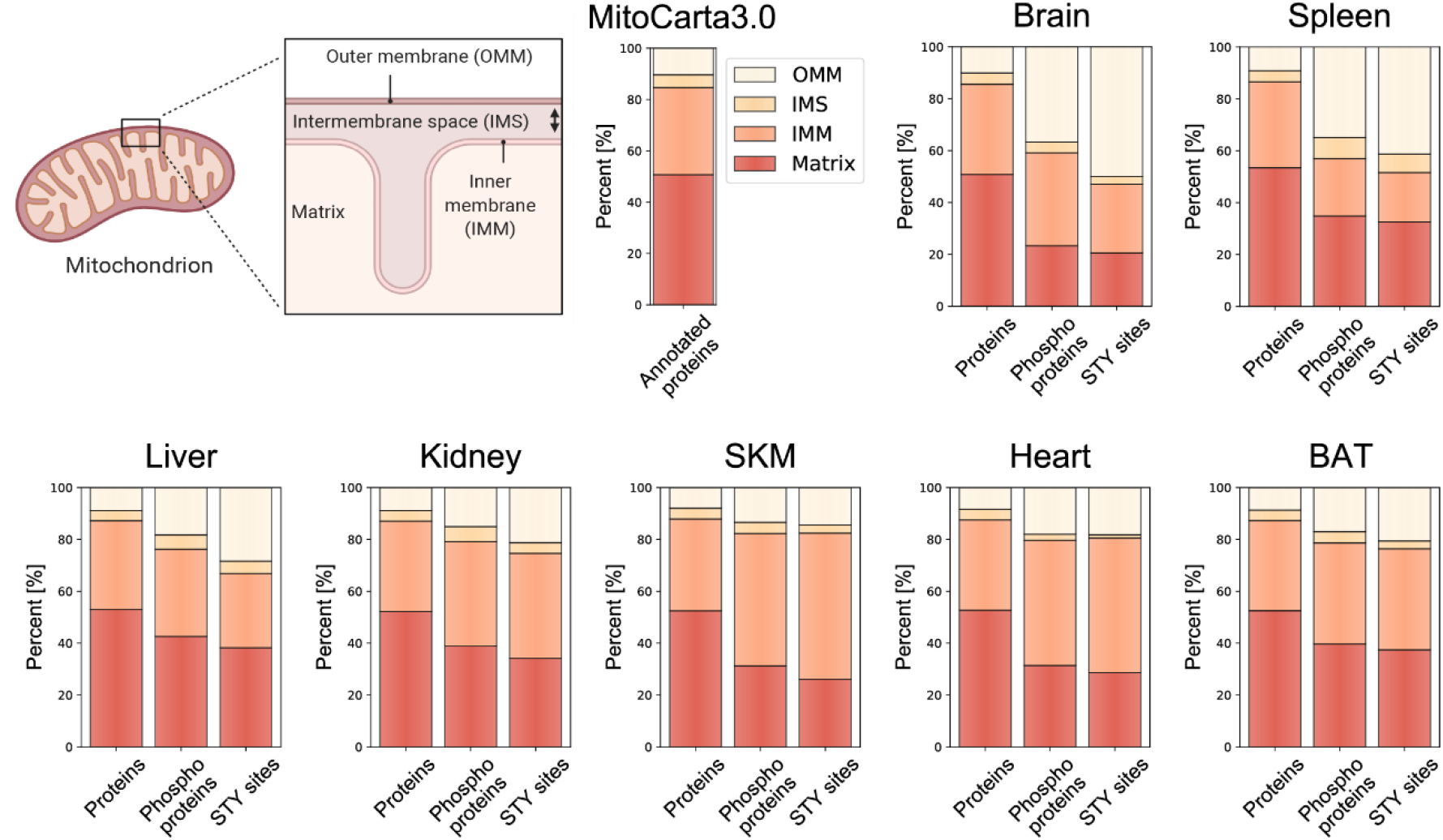
Localization distribution of mitochondrial phosphoproteome diverges from mitochondrial proteome. (A) Scheme of a mitochondrion shows four different mitochondrial localizations – outer mitochondrial membrane (OMM), intermembrane space (IMS), inner mitochondrial membrane (IMM), Matrix – and the distribution of mitochondrial proteins contained in and classified by the MitoCarta3.0 database. Bar graphs for each individual tissue show the precentral distribution of mitochondrial proteins (left), phosphoproteins (middle) and STY sites (right) across different mitochondrial localizations. Fitting of OMM proportions to a beta-regression model shows significant differences between protein-phosphoprotein and protein-STY sites for all tissues (adjusted p-value <0.0001) (see Methods and Figure 6 – Source Data 1). Skeletal muscle (SKM), brown adipose tissue (BAT).

## Supplemental information

**Supplementary Table 1.** Overview of mitochondrial phosphoproteome studies in mammalian tissue. The table was adapted and extended from Kruse at al. (Kruse and Hojlund, 2017).

| Strategy | Species | Tissue | Phosphoproteins and phosphorylation sites identified | Reference |
| --- | --- | --- | --- | --- |
| 2-DE-ProQ staining-MS | Bovine | Heart | 13 phosphoproteins | (Schulenberg et al., 2003) |
| 2-DE-ProQ staining-MS or P32 labelling-2-DE-MS | Pig | Heart | 45 phosphoproteins | (Hopper et al., 2006) |
| IMAC-MS/MS/MS | Mouse | Liver | 84 phosphorylation sites in 62 distinct proteins | (Lee et al., 2007) |
| 2D-BN-PAGE-phos—anti-Ser/Thr/Tyr Ab-MS<br>SCX/IMAC-LC-MS/MS | Rat | Heart | 2-DE: 45 phosphoproteins LC-MS/MS: 26 phosphorylation sites in 19 distinct proteins | (Feng et al., 2008) |
| Anti-Tyr Ab-MS | Rat | Brain | 7 tyrosine-phosphorylation sites in 7 distinct proteins | (Lewandrowski et al., 2008) |
| 32-P labelling/phos-tag<br>540 gel staining-2-DE-MS | Pig | Liver, Heart | 68 phosphoproteins | (Aponte et al., 2009) |
| iTRAQ labelling-SCX/HILIC/TiO2-LC-MS/MS-HCD | Pig | Heart | 56 phosphorylation sites in 38 distinct proteins | (Boja et al., 2009) |
| IMAC/TiO2-2D-nano LC-MS/MS | Rat | INS-1β cells | 84 distinct phosphoproteins | (Cui et al., 2010) |
| SAX/SCX-LC-MS/MS | Rat | Liver | 447 phosphorylation sites in 228 distinct proteins | (Deng et al., 2010) |
| SDS-PAGE-TiO2-LC-MS/MS | Mouse | Heart | 236 phosphorylation sites in 181 distinct proteins | (Deng et al., 2011) |
| SDS-PAGE-TiO2/CPP/HILIC-LC-MS/MS | Human | Muscle | 155 phosphorylation sites in 77 distinct proteins | (Zhao et al., 2011) |
| IMAC-SCX-nano-LC-MS/MS | Mouse | Liver | 811 phosphosites in 295 distinct proteins | (Grimsrud et al., 2012) |
| TiO2-HILIC-LC-MS/MS | Rat | Liver, Heart, Skeletal muscle | 899 phosphorylation sites in 354 distinct proteins | (Bak et al., 2013) |
| TiO2-HILIC-LC-MS/MS | Human | Skeletal muscle | 207 phosphorylation sites in 95 distinct proteins | (Zhao et al., 2014) |
| iTRAQ labelling-TiO2-LC-MS/MS | Human | Ovary | 124 phosphorylation sites in 67 distinct proteins | (Li et al., 2019) |
| Anti-Tyr Ab-MS | Mouse | Liver | 79 tyrosine-phosphorylated mitochondrial proteins | (Guedouari et al., 2020) |
| TMT labeling - IMAC-LC-MS/MS | Human | Skeletal muscle | 480 phosphopeptides that correspond to 196 proteins | (Sathe et al., 2021) |

